# Estrogen mediates acute elastic fibre homeostasis in skin

**DOI:** 10.1101/728865

**Authors:** Charis R Saville, David F Holmes, Joe Swift, Brian Derby, Elaine Emmerson, Matthew J Hardman, Michael J Sherratt

## Abstract

Remodelling of the dermal extracellular matrix makes a major contribution to skin fragility in the elderly. The peri-menopausal period in females is also associated with an age-like phenotype which can be reversed by hormone replacement therapy. This suggests a direct link between circulating hormone levels and tissue ageing. Despite work investigating the role of estrogen as a regulator of collagen fibril abundance and structure, the influence of estrogen on the elastic fibre system remains poorly defined. Here we used an ovariectomised (Ovx) mouse surgical menopause model to show that just 7 weeks of acute hormone deficiency significantly decreased skin tensile strength and elasticity. Systemic replacement of 17β-estradiol to physiological levels protected against these changes to the skin mechanical properties. Moreover, acute hormone deficiency differentially influenced dermal structural networks, significantly decreasing dermal elastic fibre abundance without discernible effect on collagen fibril organisation or abundance. We suggest that this specific elastic fibre proteolysis may be driven by extracellular protease activity, or be a consequence of significant adipocyte hypertrophy. 17β-estradiol supplementation in Ovx mice *in vivo* protected the elastic fibre system. Treatment of human dermal fibroblasts with 17β-estradiol *in vitro* induced the selective upregulation of tropoelastin, fibrillin-1 and associated elastic fibre-associated proteins (including EMILINs and fibulins). In summary, these data show that the elastic fibre system is significantly perturbed by estrogen deprivation. Thus, pharmacological intervention may slow the acute effects of menopause and potentially the chronic effects of ageing in skin.

## INTRODUCTION

The onset of menopause and the subsequent dramatic loss of circulating sex steroid hormones, particularly 17β-estradiol, has profound adverse effects on tissue homeostasis and function in disparate organs systems, including the blood vessels, lungs, bone and skin [1–4]. In the skin, age-associated estrogen deprivation has been shown to be an important regulator of delayed wound healing affecting a greater number of wound associated genes than age alone, highlighting the importance of estrogen in regulating timely cutaneous would healing [5]. Aside from wound healing, in intact tissues both ageing and estrogen-deprivation are associated with compositional and structural remodelling of the dermal extracellular matrix (ECM), with consequential loss of tissue resilience and increased tissue fragility [6–8]. These functional changes are thought to directly increase vulnerability of the elderly population to skin damage resulting from friction and pressure, which severely impact quality of life and healthcare costs [9].

As yet, the pathological mechanisms which drive this menopause-associated loss of mechanical fidelity are poorly defined. However, the ECM-rich skin dermis, and in particular the most abundant dermal proteins, type-I collagens, are considered to be the dominant mediators of skin mechanics [10]. Collagen-I, forms a “basket-weave” arrangement of tensile strength-conferring fibril bundles [11, 12]. These fibrils are lost, or structurally remodelled as a consequence of estrogen deprivation; menopause [13–16] or ovariectomy (Ovx) in animal models [17–19]. By contrast, 17β-estradiol supplementation promotes collagen deposition [14, 15, 20]. Surgical Ovx in rodents models many aspects of human menopause including osteoporosis, neurodegeneration and cardiac dysfunction [21–23], impaired elastic recoil and accelerated photo-damage in skin [4, 24, 25]. In the sheep, Ovx has been associated with long-term remodelling of collagen fibril structure in skin and bone [24, 26].

Fibrillar collagens are not the only fibrous dermal component. Elastic fibres, although less abundant than collagen fibrils, play key roles in mediating mechanical resilience (passive recoil) and tissue phenotype via the sequestration of growth factors [27, 28]. They are comprised of a cross-linked elastin core surrounded by an outer mantle of fibrillin microfibrils [29] which in turn binds proteins including microfibrillar-associated protein 2 (MFAP2, also known as MAGP-1), and members of the elastin microfibril interface-located protein (EMILIN) and fibulin families [29–31]. Individual elastic fibre components are particularly sensitive to oxidative damage arising from a wide range of biological and environmental factors, such as ultra violet radiation exposure, smoking, diabetes and/or raised body mass index (BMI) [32–37]. Estrogen acts as a direct anti-oxidant, inducing the expression of antioxidant enzymes and mediating hypertrophy of sub-cutaneous fat [38–40]. However, the influence of circulating estrogen on elastic fibre biology remains poorly understood. In this study, we have combined murine *in vivo* and human *in vitro* studies to test the hypothesis that estrogen directly mediates elastic fibre proteostasis in skin.

## MATERIALS AND METHODS

### Reagents, tissue samples and cultured cells

All reagents were obtained from Sigma-Aldrich Co. Ltd (Poole, UK) or BDH Ltd (Poole, UK), unless otherwise specified. Procedures involving mice accorded to the UK Animals (Scientific Procedures) Act 1986 under UK home office licence (40/3713). All mice used in this study were wild type C57BL/6 females (Envigo, UK) aged 7 weeks at the start of the experimental period. Primary human dermal fibroblasts (HDFs) were derived from female abdominal skin and maintained in phenol red-free DMEM supplemented with L-glutamine, 10% charcoal-stripped foetal calf serum and penicillin-streptomycin.

### Ovariectomy and estrogen supplementation

Estrogen deprivation was induced in five mice (Ovx group) by bilateral ovariectomy as previously described [41]. Mice were estrogen-deprived for a period of seven weeks. Briefly, the mice were anaesthetised and following ventral laparotomy, the ovaries were removed using sterile scissors. The body wall and skin were closed using sutures and buprenorphine (0.1 mg/kg) administered as analgesia. Estrogen deprivation was confirmed by uterine atrophy. The potential protective effects of exogenous estrogen in skin were characterised in five Ovx mice which were also treated with a 60 day release subcutaneous 17β-estradiol replacement pellet (0.1 mg) (Innovative research of America, Florida, USA) inserted at the base of the neck (Ovx+E group). A final group of five mice served as age-matched controls (Intact group). This was repeated on a second group of mice which were estrogen-deprived for 3 weeks prior to collection.

### Mechanical characterisation of murine skin

The mechanical effects of Ovx and 17β-estradiol supplementation were determined by stretching 10 mm wide x 30 mm long strips of ventral skin from intact control, Ovx and Ovx+E (*n* = 5) mice using an Instron 3344 100 N load cell (Instron, USA). The tissue was loaded to failure at a rate of 20 mm/min [42]. Skin viscoelasticity in intact control and Ovx mice (*n* = 5) was characterised by testing the stress relaxation behaviour of ventral skin in a PBS bath using an Instron 5943 10N load cell (Instron, USA). Skin was preconditioned by cyclical loading to 1 N at a rate of 10 mm/min (5 repeats) and then loaded to 1 N and held at a constant strain for 160 seconds. All data was collected and analysed using Bluehill software (Instron, USA).

### Histological characterisation of murine skin: collagen and elastic fibre abundance, organisation and depth of subcutaneous fat

Histological sections (6 µm) were prepared from formalin-fixed, paraffin-embedded dorsal skin of intact control, Ovx and Ovx+E (*n* = 5) mice. Elastic fibres were stained using Gomori’s aldehyde fuchsin [43] and visualised by bright-field optical microscopy (Nikon eclipse E600 microscope/SPOT camera). Total tissue collagen was assessed by staining with Masson’s trichrome which is reported to stain amorphous collagen [44]. The abundance of organised fibrillar collagen was assessed by measuring collagen birefringence following staining with picrosirius red (PSR) and imaging by polarised light (Leica DMRB) [45]. The proportion of tissue area occupied by elastic fibres, amorphous collagen and organised fibrillar collagen was semi-quantitatively assessed using ImageJ 1.46r software as previously described [45, 46]. The ratio of thick and thin collagen fibrils was semi-quantitatively measured using the colour deconvolution plugin to split colour channels, a threshold applied and measured as percentage area [42] (Image J, National Institutes of Health, USA). Finally, the collagen orientation (coherency) in the PSR/polarised light images was assessed as previously described [45, 47].

### In situ gelatinase activity in cryo-preserved skin sections

In order to quantify and localise the effects of Ovx and Ovx+E on mouse skin gelatinase activity we used *in situ* gelatin zymography [37]. Cryo-preserved skin in optimal cutting temperature (OCT) compound was sectioned to a nominal thickness of 10 µm (Leica CM3050 cryostat). DQ gelatin (Sigma, UK) solution (1 mg/ml DQ gelatin in low gelling temperature agarose) containing DAPI (1 μg/ml) was pipetted over the section and covered with a glass coverslip. Sections were incubated for 18 hours at 4 °C and images captured using an Olympus BX51 microscope with a CoolSnapES camera and Metavue software (Molecular Devices). Fluorescence was quantified as previously described using ImageJ software (version 1.46r; National Institutes of Health, USA) [37].

### In vivo expression of murine elastic fibre genes

RNA was extracted from snap frozen skin to assess expression of elastic fibre components tropoelastin (*Eln*) and fibrillin-1 (*Fbn1*) following a standard protocol (Life Technologies, UK). Briefly tissue was homogenised in TRIzol reagent and extracted in chloroform then purified using the RNA Purelink kit (Life Technologies, UK). Subsequently, cDNA was transcribed from RNA (Reverse Transcriptase kit, Promega, Madison, USA) using Reverse Transcriptase (Roche, UK). Quantitative PCR was carried out on an MyiQ thermal cycler (Bio-Rad, UK) with cDNA diluted over three orders of magnitude and expression ratios normalised to housekeeping genes *Gapdh* and *Ywhaz*.

### ECM gene expression by cultured human cells

Human keratinocytes and HDFs were cultured overnight in 12 well plates (seeding density 0.17 × 10^6^ cells per well) in a CO_2_ incubator at 37 °C. 17β-estradiol was added to a final concentration of 1 μM, and cells were harvested 6 or 24 hours post-treatment by removal of the media and addition of TRIzol reagent (Life Technologies, UK). RNA was extracted using the RNA Purelink kit (Life Technologies, UK), cDNA synthesis and quantitative PCR were carried out as described above for *FBN1* and housekeeping genes *GAPDH* and *YWAHZ*.

### Protein synthesis by cultured human fibroblasts: mass spectrometry (MS)

Reagents for MS were purchased from Fisher Scientific (UK), unless otherwise noted. Primary HDFs were plated into 2 x 6 well plates at a seeding density of 50% confluence and allowed to adhere overnight. Fresh media was added on alternate days and half of the samples (6 wells) were supplemented with 17β-estradiol (1 μM final concentration). After eight days of culture, ECM proteins were removed from the dish by washing for 20 minutes in a minimal volume of high-salt extraction buffer (2 M NaCl, 25 mM ammonium bicarbonate, 25 mM dithiothretol). Extracted proteins were digested overnight using immobilized-trypsin beads (Perfinity Biosciences) in 1 mM CaCl_2_ and 25 mM ammonium bicarbonate, prior to reduction (10 mM dithiothretol) and alkylation (30 mM iodoacetamide). The samples were acidified with 0.4% trifluoracetic acid, cleaned by biphasic extraction (vortexing with ethyl acetate) and dried down in a speed-vac. Peptides were desalted using POROS R3 beads according to the manufacturer’s protocol (Thermo Fisher). Samples were dried down in a speed-vac, dissolved in injection buffer (5% HPLC-grade acetonitrile and 0.1% trifluoroacetic acid in deionised water) and stored at 4 °C prior to analysis by mass spectrometry.

The digested samples were analysed by liquid chromatography tandem MS (LC-MS/MS) using an UltiMate 3000 Rapid Separation liquid chromatography system (RSLC, Dionex Corporation, Sunnyvale, CA) coupled to an Orbitrap Elite (Thermo Fisher Scientific, Waltham, MA) mass spectrometer. Peptide mixtures were separated using a gradient from 92% A (0.1% FA in water) and 8% B (0.1% FA in acetonitrile) to 33% B, in 104 min at 300 nL min^−1^, using a 75 mm x 250 μm inner diameter 1.7 μM CSH C18, analytical column (Waters). Peptides were selected for fragmentation automatically by data dependant analysis. MS spectra from multiple samples were aligned using Progenesis QI (Nonlinear Dynamics) and searched using Mascot (Matrix Science UK), against the SWISS-Prot and TREMBL human databases. The peptide database was modified to search for alkylated cysteine residues (monoisotopic mass change, 57.021 Da), oxidized methionine (15.995 Da), hydroxylation of asparagine, aspartic acid, proline or lysine (15.995 Da) and phosphorylation of serine, tyrosine, threonine, histidine or aspartate (79.966 Da). A maximum of two missed cleavages was allowed. Peptide detection intensities were exported from Progenesis QI as Excel spreadsheets (Microsoft) for further processing. Peptide signal intensities were normalised to the geometric mean of all ECM components (defined by gene ontology annotation in the UniProt database). Fold changes in ECM protein levels were then calculated from changes to median peptide signal intensities.

### Protein synthesis by cultured human fibroblasts: western blotting

HDFs (*n* = 3) were treated with 17β-estradiol supplemented (1 μM) or control media on days 1, 3, 7 and 10. After 14 days, media was collected and added to Laemmli sample buffer (Bio-Rad) containing 5% 2-mercaptoethanol (Bio-Rad), electrophoresed by SDS-PAGE and blotted onto nitrocellulose membrane. Membranes were blocked with 5% non-fat milk 0.1% tween (Sigma, UK) prior to incubation with primary antibodies to elastin (Santa Cruz SC-17580) used at 1:200 dilution or β-actin (Sigma A5441) used at 1:5000 dilution and detection with peroxidase-labelled secondary antibodies (GE Healthcare, UK) and ECL Plus (enhanced chemiluminescence; GE Healthcare, UK).

## RESULTS

### Estrogen modulates skin tensile strength, stiffness and viscoelasticity

We first assessed the effects of acute estrogen deprivation, and the protective effects of 17β-estradiol supplementation, on the skin functional parameters of tensile strength and stiffness. For strips of ventral skin loaded to failure at a uniform extension rate (Fig. 1A), breaking stress (and therefore tensile strength) was significantly lower in Ovx mouse skin compared to control skin (42% reduction; *p* < 0.05). 17β-estradiol replacement (Ovx+E) fully restored breaking stress to pre-Ovx levels (Fig. 1B). Skin stiffness (Young’s modulus) was also significantly reduced in Ovx compared to intact control (45% reduction; *p* < 0.05) and almost entirely preserved in Ovx+E skin (Fig. 1C). The strain required to break the skin was not significantly altered in Ovx or Ovx+E samples (Fig. 1D). Collectively, these data demonstrate that acute (7 weeks) estrogen deprivation profoundly affects skin tensile strength and stiffness.

**Figure 1.**
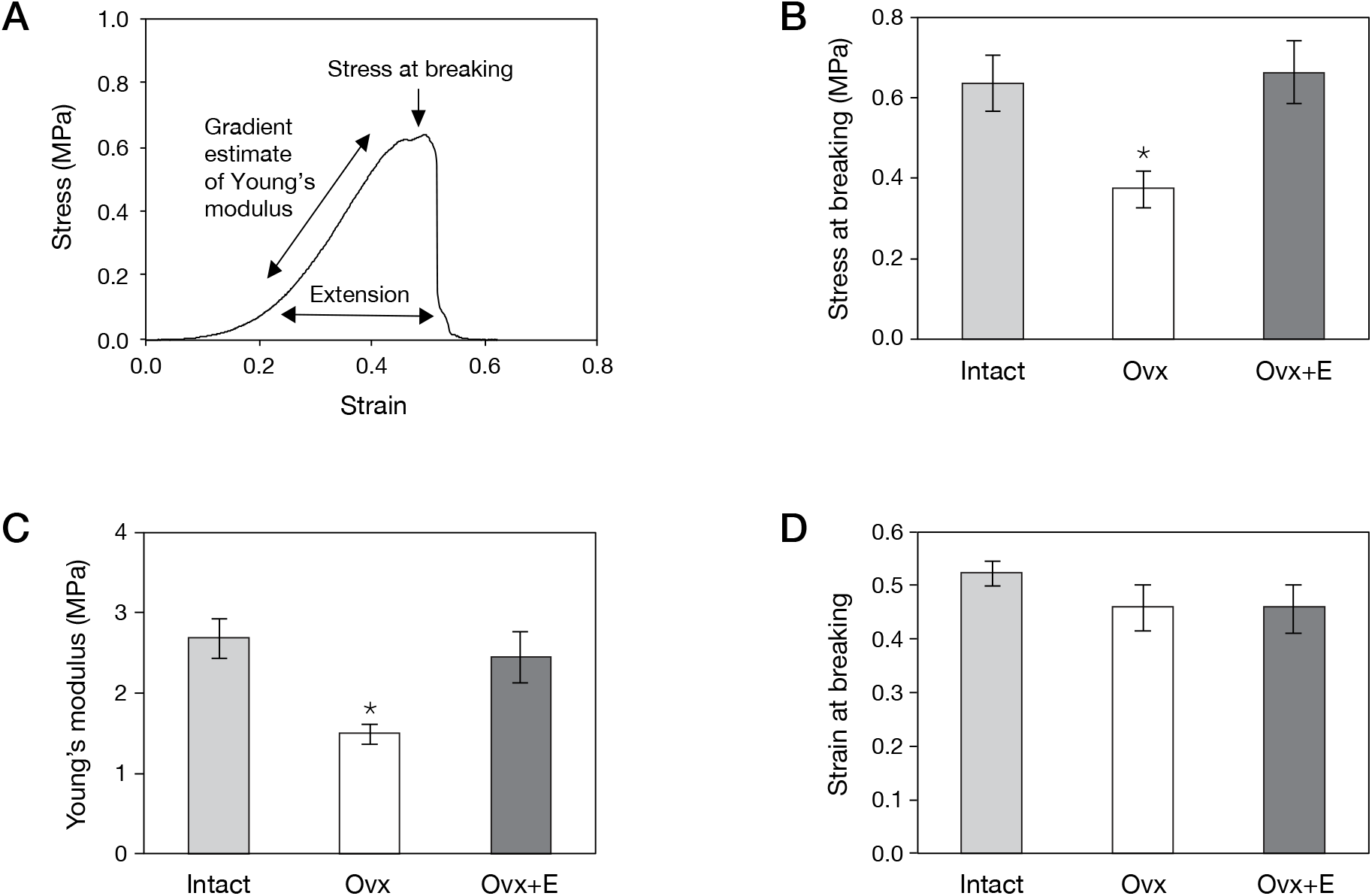
Estrogen modulates the mechanical properties of murine skin. (**A**) Stress:strain curve permits estimate of the Young’s modulus (gradient of the linear section), tensile strength and strain (extension) at breaking of ventral skin strips (1 cm wide). (**B**) The tensile strength of skin was reduced following seven weeks of estrogen deprivation (Ovx), a change fully protected against with 17β-estradiol treatment (Ovx+E). (**C**) The Young’s modulus was also significantly reduced in Ovx compared to Intact and Ovx+E skin. (**D**) The strain at breaking was not significantly affected by Ovx or Ovx+E treatments. In plots (B)-(D), bars indicate mean ± s.e.m. N ≥ 4; * *p* < 0.02.

### Dermal collagen abundance and organisation are insensitive to estrogen modulation

As chronic estrogen deprivation has previously been linked to changes in both skin mechanics and dermal collagens [13, 14, 16, 20] we used histological approaches to quantify the area fractions of amorphous and organised fibrillar collagen in our acute estrogen deficiency and rescue models. Surprisingly, the mechanical alteration observed following Ovx (Fig. 1) was not correlated with a change in total collagen content as measured using Masson’s trichrome (Figs. 2A and B). In order to assay the effects of estrogen on mechanically robust fibrils and fibril bundles [12, 48] we measured birefringence in collagen fibrils stained with picrosirius red (PSR). The presence of organised fibril bundles has been associated with local mechanical stiffening in breast stroma [45] but we did not observe any relationship between collagen birefringence and circulating estrogen (Figs. 2C-F). Therefore, we conclude that mechanical changes due to acute (7 week) estrogen deprivation do not appear to be mediated by fibrillar collagen remodelling.

**Figure 2.**
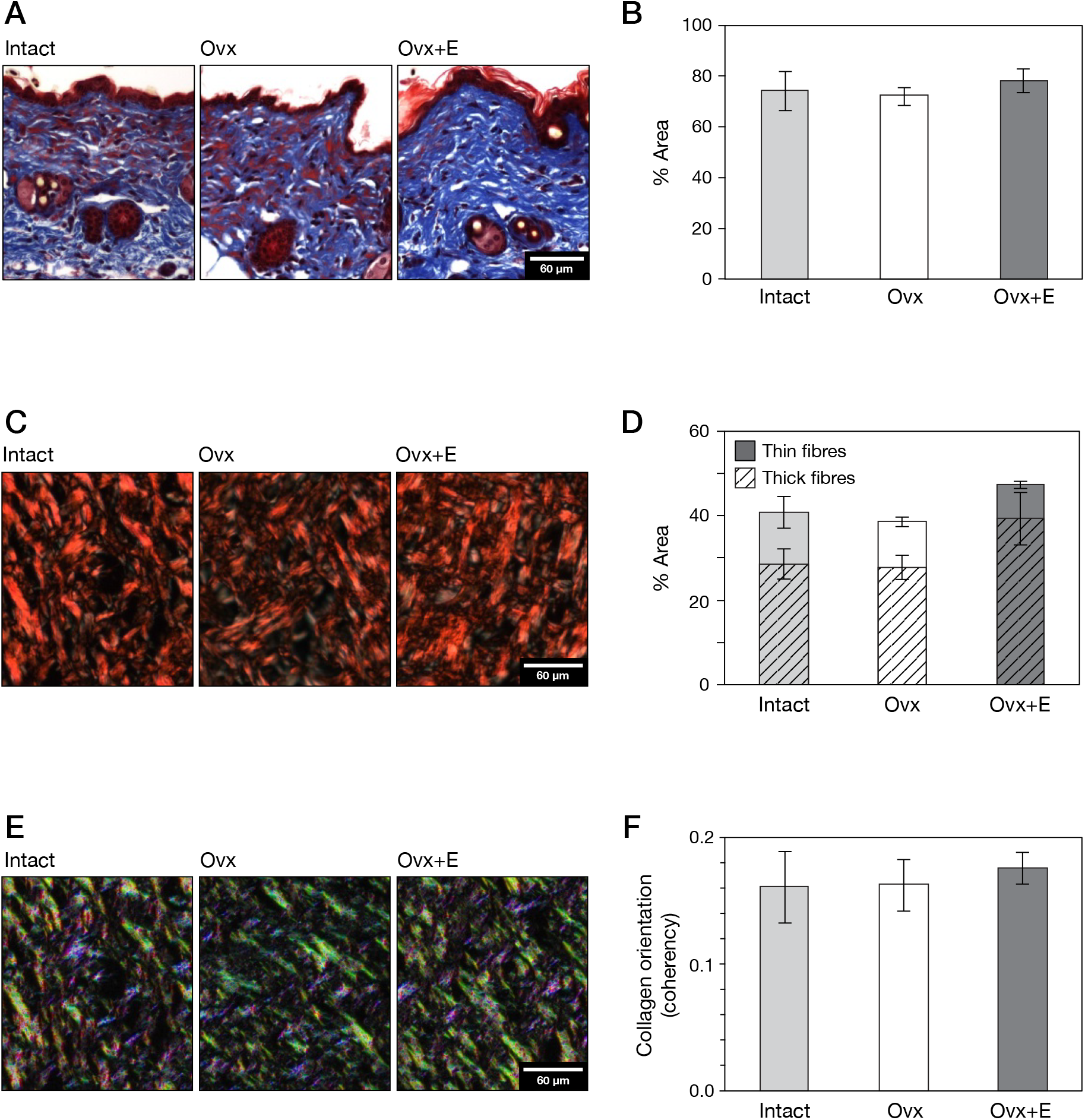
Dermal abundance and organisation is insensitive to circulating estrogen. (**A**) Sections of mouse skin from control (Intact), estrogen deprived (Ovx) and 17β-estradiol supplemented (Ovx+E) treatment groups, stained against total collagen with Masson’s trichrome stain. (**B**) Total collagen was unaltered following Ovx or 17β-estradiol replacement. (**C**) Picrosirius red staining imaged under polarised light allowed visualisation of fibrillar collagen. (**D**) There was no effect on fibrillar collagen following Ovx or 17β-estradiol replacement. (**E**) Collagen fibril orientation was analysed using Orientation J software. (**F**) Quantification of collagen fibril orientation again showed no inter-group differences. In plots (B), (D) and (F), bars indicate mean ± s.e.m. N = 5.

### Estrogen deficiency affects both elastic fibre abundance and fibrillin-1 expression *in vivo*

Estrogen deprivation altered elastic fibre morphology: compared with both intact control and Ovx+E mice, elastic fibre positive material in Ovx skin was punctate rather than fibrillar (Fig. 3A). This estrogen-associated loss of fibrillar architecture was accompanied by a significant (37%; *p* < 0.01) reduction in tissue area occupied by elastic fibres, and effect that was entirely reversed by 17β-estradiol supplementation (Figs. 3A and B). Fibrillin-1, but not tropoelastin, expression was also significantly reduced in the skin of Ovx mice (Figs. 3C and D). Conversely, fibrillin-1 but not tropoelastin expression was strongly elevated in the skin of Ovx+E mice compared to the Ovx group (Figs. 3C and D). As these observations suggest that estrogen may mediate both elastic fibre synthesis and degradation we next investigated the effects of Ovx on gelatinase activity using gelatin zymography. Further to this, a suggested link between increased matrix metalloproteinase (MMP) activity and increased adipocyte size led us to also investigate the sub-cutaneous adipocytes in this model [34].

**Figure 3.**
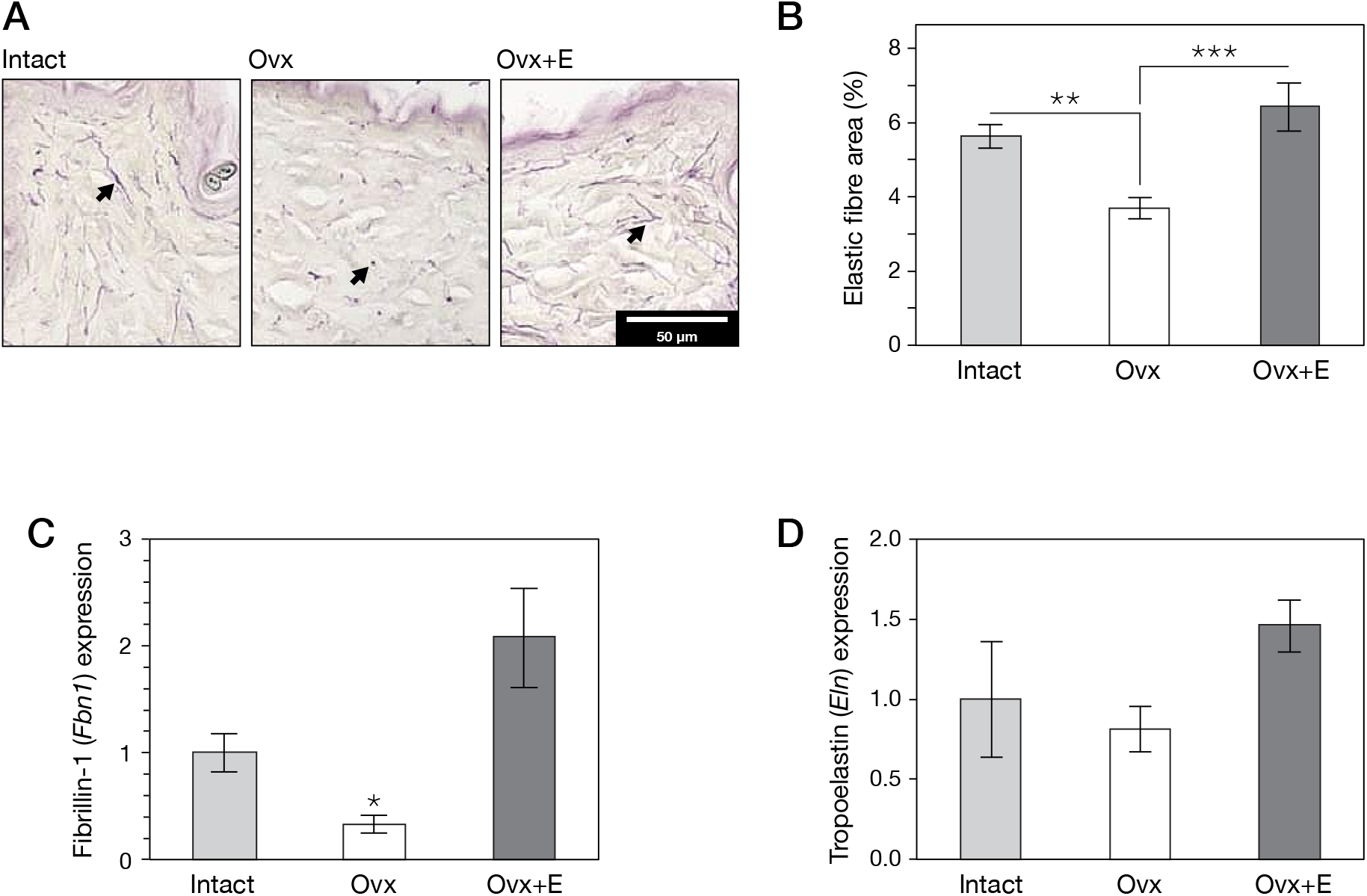
Estrogen mediates elastic fibre degradation and synthesis. (**A**) Intact, Ovx and Ovx+E skin sections with Gomori’s aldehyde fuchsin staining against tissue elastic fibres (dark purple). The elongated elastic fibres in Intact and Ovx+E skin were replaced by punctate staining in Ovx mice (arrows). (**B**) Both intact control and Ovx+E skin contained significantly more elastic fibre positive material than Ovx skin. Elastic fibres occupied a smaller area (5 – 6%) than fibrillar collagens in the Intact control. (**C**) Fibrillin-1 transcript (*Fbn1*), relative to housekeeping genes and normalised to the Intact sample. (**D**) Tropoelastin transcript (*Eln*), relative to housekeeping genes and normalised to the Intact sample. Plots show mean ± s.e.m. N ≥ 4; *** *p* < 0.001 ** *p* < 0.01 * *p* < 0.05.

### Estrogen loss increases gelatinase activity, and adipocyte hypertrophy and proliferation

ECM proteins are primarily degraded by either MMPs or cathepsins [49]. Here we used an *in situ* gelatin zymography approach to localise and quantify relative gelatinase activity in cryo-preserved murine skin sections. Using this assay proteolytic digestion of the fluorescein labelled gelatin led to florescence, shown here as bright white [50]. Dermal gelatinase activity was increased by three weeks post-Ovx compared with intact control (Figs. 4A and B). Again, we observed that estrogen supplementation was protective. Increased gelatinase activity in estrogen-deprived dermis could be driven by dermal fibroblasts [51], epidermal keratinocytes [52], or hypodermal adipocytes [41]. Although the epidermis was clearly a highly gelantinolytic environment, gelatinase activity in the Ovx group was observed throughout the dermis. Given the known association between increased subcutaneous fat, adipokine-mediated local protease activity and elastic fibre remodelling, we characterised the effects of Ovx and Ovx+E on the hypodermis. Ovx induced a significant increase in the depth of the hypodermis and the size of individual adipocytes, which was reversed by 17β-estradiol (Figs. 4C-E).

**Figure 4.**
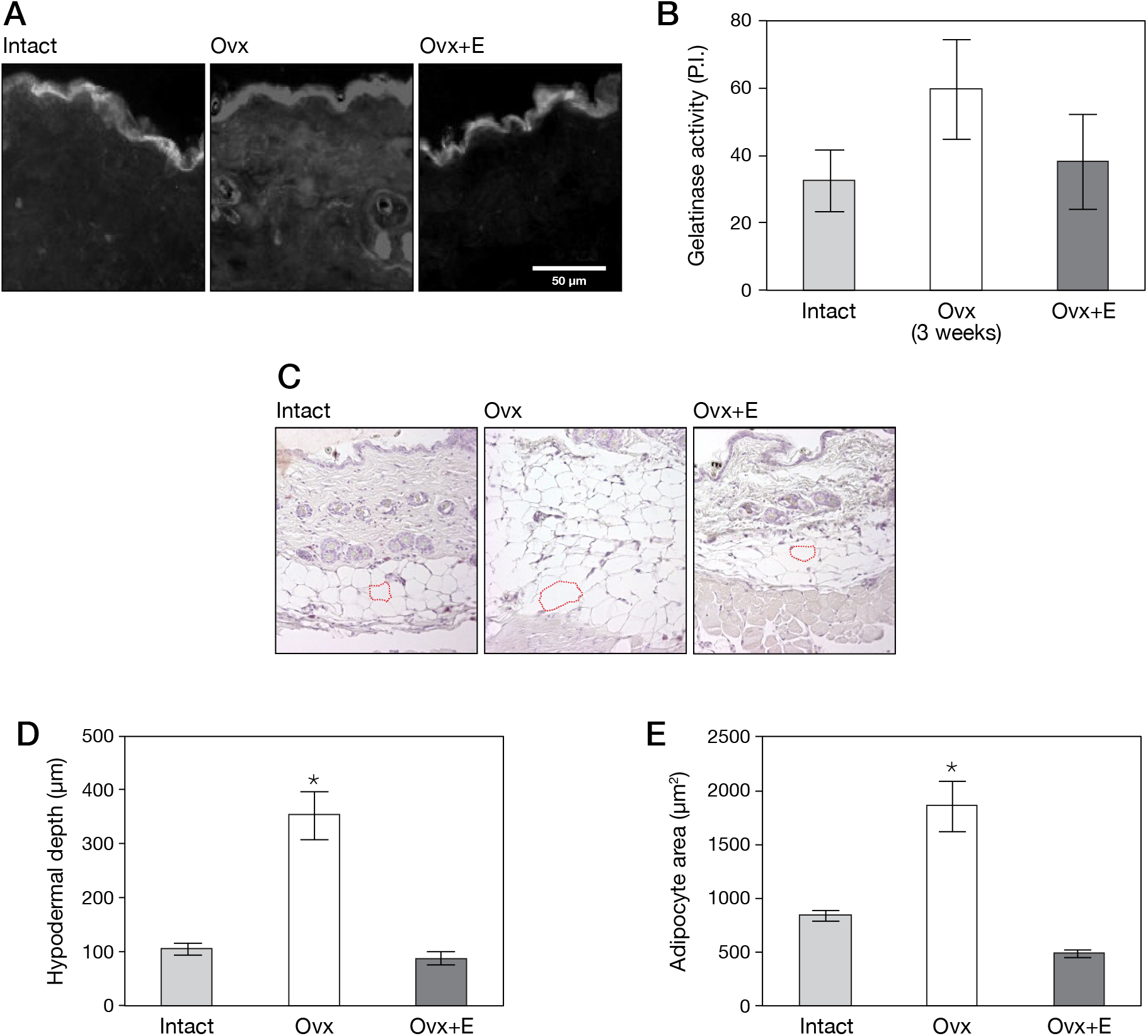
Loss of circulating estrogen induces gelatinase activity and hypodermal hypertrophy. (**A**) *In situ* gelatin zymography of murine skin three weeks post Ovx and Ovx+E. White areas indicate gelatinase activity. (**B**) Quantification of gelatinase activity showed an increase three weeks post Ovx, although this effect was not significant. (**C**) Haematoxylin and eosin stained sections of murine skin. Individual adipocytes are indicated in red. (**D**) Analysis of the stained sections showed that hypodermal depth was significantly increased in the Ovx group. (**E**) The size of adipocytes was also significantly increased in the Ovx group. Plots show mean ± s.e.m. N = 5; * *p* < 0.0001.

### Estrogen mediates *in vitro* expression of elastic fibre associated proteins by dermal fibroblasts

Given that our murine data indicate major effects of estrogen on skin elastic fibres, we next looked for cross-species conservation, exploring the effect of estrogen on synthesis of ECM proteins by cultured human skin cells. Firstly, we treated primary human keratinocytes and HDFs with 17β-estradiol *in vitro*. In each case 17β-estradiol treatment led to rapid, statistically significant up-regulation of fibrillin-1 (FBN1) mRNA (Figs. 5A and B). To explore the longer-term effects of 17β-estradiol we turned to proteomic analysis of cell-derived matrix by mass spectrometry (Fig. 5C). Five proteins associated with elastogenesis (EMILIN1, EMILIN2, FBN1, FBLN2 and FBLN5) were upregulated, while MMPs 1, 2 and 3 were downregulated. One key elastic fibre component missing from this analysis was elastin, presumably because elastin is highly crosslinked and therefore extremely difficult to detect by mass spectrometry. As an alternative strategy we used western blotting of cell media to detect the soluble precursor tropoelastin (Fig. 5D). Here, tropoelastin was significantly elevated in HDF cell media following 17β-estradiol treatment.

**Figure 5.**
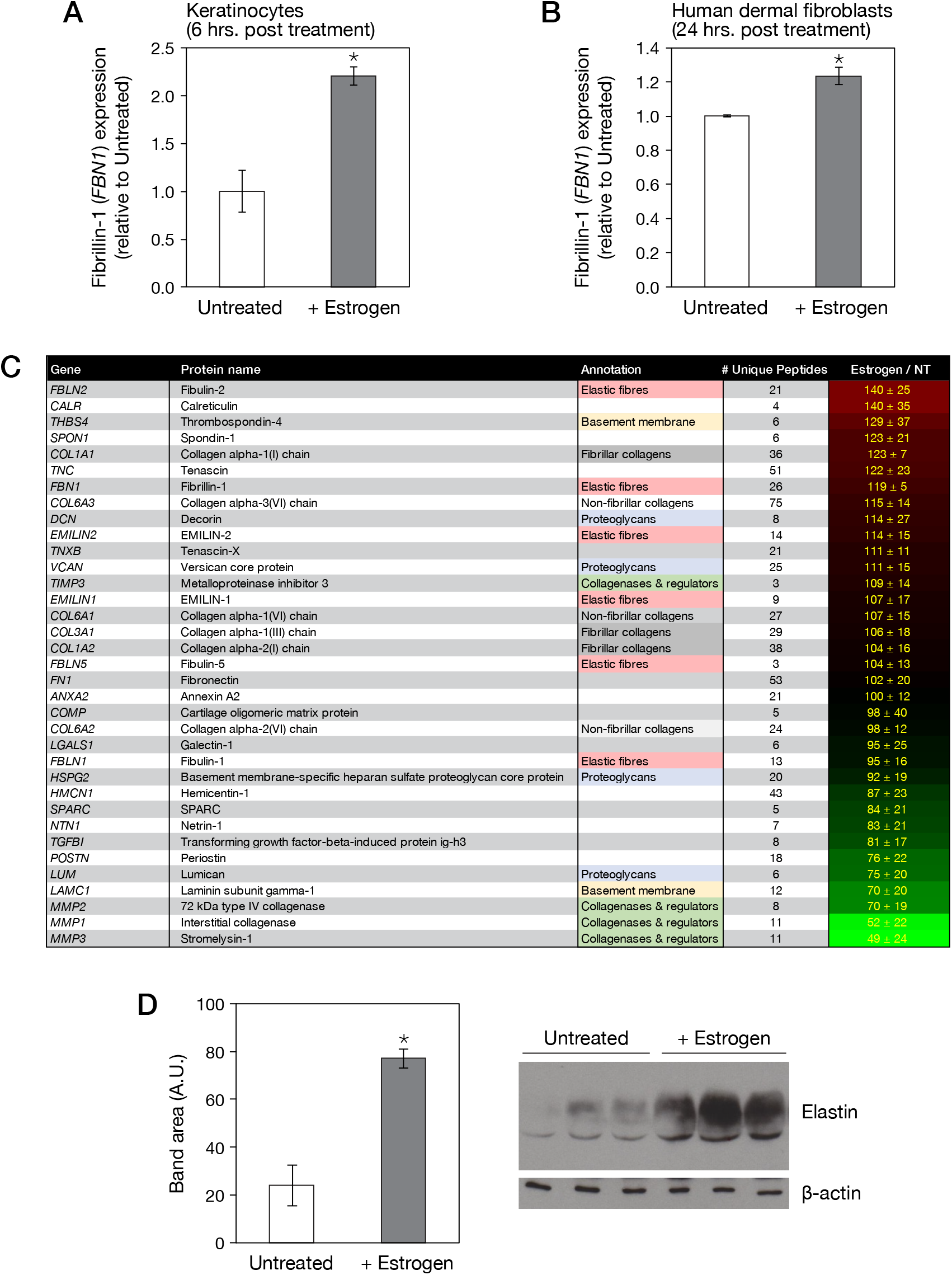
Estrogen mediates the expression of elastic fibre proteins. (**A**) Quantification of fibrillin-1 transcript (*FBN1*) in human keratinocytes, with and without 17β-estradiol treatment (+Estrogen), normalised to untreated control cells and housekeeping genes. (**B**) Quantification of *FBN1* in human dermal fibroblasts (HDFs), relative to control cells and housekeeping genes. Both cells types increase fibrillin-1 expression in response to 17β-estradiol treatment. (**C**) Mass spectrometry quantification of extracellular matrix (ECM; identified by gene ontology annotation) proteins secreted by HDFs cultured for eight days. The effects of estrogen treatment on ECM production is expressed as a percentage relative to untreated controls ± s.e.m (N = 3). Estrogen upregulated a number of elastic fibre associated components, but decreased levels of MMPs. (**D**) Western blot analysis showed that HDF-derived tropoelastin was significantly upregulated by 17β-estradiol treatment. Plots (A), (B) and (D) show mean ± s.e.m. N = 3; * *p* < 0.01.

## DISCUSSION

In this study and earlier work [16] we have shown that acute (seven week) estrogen deprivation had a major impact on the mechanical properties of murine skin, leaving the organ weaker, more compliant and easily deformed. These mechanical changes were associated with significant remodelling of dermal elastic fibres, adipocyte hypertrophy and increased gelatinase activity. Supplementation with exogenous 17β-estradiol reversed these *in vivo* changes and directly upregulated the expression of elastic fibre components by cultured human cutaneous cells *in vitro* (Fig. 6).

**Figure 6.**
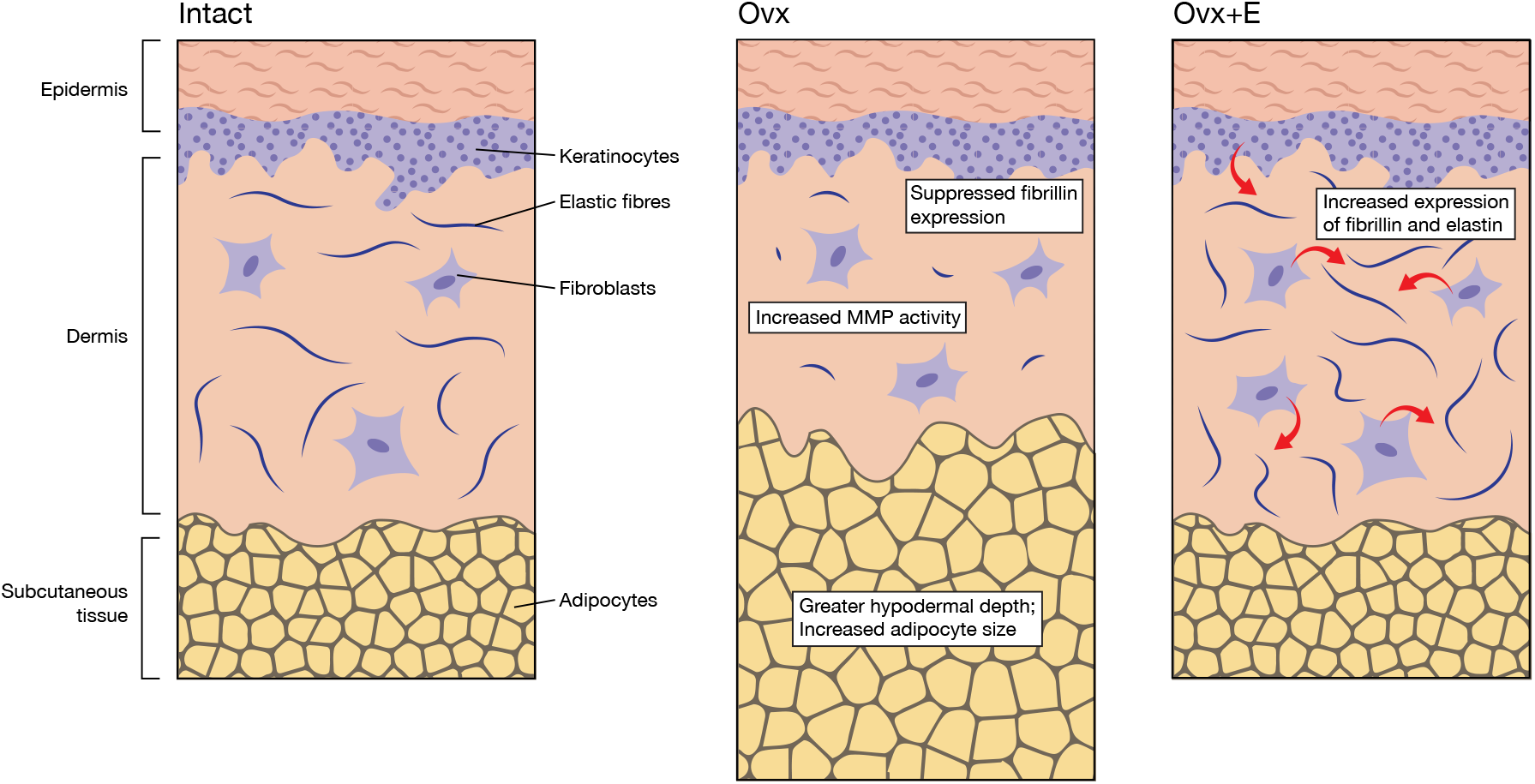
Schematic representation of skin following estrogen deprivation and replacement. Estrogen deprivation leads to an increase in subcutaneous adipose tissue thickness and local MMP activity, causing elastic fibre degradation and suppression of fibrillin production. Estrogen treatment upregulates fibrillin expression in both fibroblasts and keratinocytes, induces tropoelastin in fibroblasts alone, and dampens local MMPs.

*In vivo*, estrogen deficiency substantially reduced the tensile strength, stiffness and resilience of murine skin. Loss of elastic recoil (manifesting as a prolonged skin tenting after pinching) has previously been described in Ovx mice [4]. A longer-term study in rats also reported an Ovx induced loss of tensile strength when measured using a tensometer [53]. In human skin Pierard *et al.* report increased skin extensibility in the peri-menopausal period [54] which aligns with our observations of reduced Young’s modulus in Ovx mice. There remains surprisingly little data on the mechanics of post-menopausal skin and studies of human skin variously report that Young’s modulus increases [55], decreases [56] and remains unchanged with age. The skin, like many biological tissues, is structurally complex displaying material properties that are viscoelastic [57]. Ovariectomy has been shown to alter the stress relaxation response, accelerating the rate at which skin relaxes versus intact control mice, highlighting that estrogen-deprived skin becomes lax under sustained force. Collectively these mechanical findings suggest that estrogen deficiency leads to an acute remodelling of the cutaneous ECM which mirrors the increased laxity and weakness of aged human skin. This could be a key contributing factor in the increased susceptibility of aged or indeed estrogen-deprived skin to developing chronic wounds, a problem further compounded by the direct role of estrogen in timely wound healing [58].

Age-associated changes in the appearance of skin have been attributed to a loss of fibrillar collagen [8, 59], with inferred effects on tensile strength [60]. Chronic (greater than two years) estrogen deprivation in an ovine Ovx model was associated with dermal collagen remodelling [24]. By contrast, acute (seven week) estrogen deprivation in our Ovx mouse model had no observable effect on fibrillar collagen. Instead there was a significant loss of elastic fibres accompanied by reduced expression of the key elastic fibre component *fibrillin-1*. The rapidity of this loss was surprising, as elastic fibres are thought to be subject to very little turnover [32, 61, 62]. Our data suggest that elastic fibre remodelling may be particularly important in early skin changes following menopause.

The mechanism(s) by which elastic fibres are remodelled in acute estrogen deprivation remain to be elucidated. The process is likely to be multifactorial with an altered balance between degradation and synthesis. A recent study by Ezure *et al.* draws a clear link between raised BMI, elastic fibre degradation and MMP expression [34]. As increased adiposity was also evident in our estrogen deprived murine skin, we localised cutaneous gelatinase activity in three-weeks post Ovx mice. The observation of raised gelatinase activity (which may be attributable to MMPs [63]) implied that elastic fibre remodelling in acute Ovx may be enzyme driven. Mass spectrometry data generated when HDFs were treated with estrogen showed a reduction in MMP expression, confirming estrogen has roles in suppressing MMP activity. Alternatively, adipocyte hypertrophy has been associated with increased oxidative stress [64]. We have previously shown that fibrillin-1 and other EGF-based elastic-fibre-associated proteins are enriched in oxidatively sensitive Cys, Trp, Tyr and Met residues [65] and have suggested that these proteins act as endogenous extracellular antioxidants [33]. Estrogen itself may protect the tissues and elastic fibres both via its intrinsic anti-oxidative properties and the induction of antioxidant enzymes [66].

Our *in vivo* data are also consistent with estrogen directly inducing synthesis of new elastic fibres. This scenario clearly challenges the current dogma that elastic fibres are only laid down during development (Reviewed in [32]). To test this observation we used mass spectrometry to quantitatively analyse the matrix deposited *in vitro* from isolated human HDFs treated directly with 17β-estradiol. We found numerous ECM proteins were induced or repressed following estrogen treatment, with four identified proteins specifically linked to elastic fibre deposition. Fibrillin-1 provides the initial scaffold onto which elastin localises as the elastic fibre builds [29]. EMILIN-1 localises to the microfibril-elastin interface and may play important roles in the regulating tropoelastin deposition [30]. We also found induction of EMILIN-2, a less well-characterised family member which shares 70% similarity to EMILIN-1 [67]. Intriguingly, EMILIN-1 and EMILIN-2 can interact in a head to tail arrangement to form larger assemblies [68]. Thus, the co-ordinated induction of both family members by estrogen may be important in the context of elastic fibre assembly. Fibulin-2 is also found at the elastin microfibril interface and in skin is found to co-localise with fibrillin-1 [69], suggesting an important role in the synthesis of mature elastic fibres. Finally, tropoelastin was strongly upregulated in human dermal fibroblasts following 17β-estradiol treatment, further confirming the role of estrogen in elastic fibre remodelling. These observations suggest that estrogen acts as a potent inducer of cutaneous elastic fibre synthesis in both mouse and human. These data are supported by other human studies reporting increased mRNA for both fibrillin-1 and tropoelastin [70] or an increase in total elastic fibres [71] following topical estrogen treatment. In addition to their role in skin, elastic fibres are critical for normal function of respiratory and cardiovascular tissues. In these systems rapid loss of elastic fibres following menopause would contribute to serious health issues. It is therefore likely that compensatory mechanisms exist. For example, estrogen receptors (ERs) can signal in a ligand independent manner via growth factors such as insulin-like growth factor (IGF) [41, 72, 73]. This ligand independent signalling may become more prevalent in the postmenopausal period. The skin also contains all the enzymes required to synthesise steroid hormones from cholesterol and this provides a major source of estrone in postmenopausal skin [74]. Indeed, a more detailed understanding of estrogen’s dramatic effects on the elastic fibre system is essential to develop new pharmacological interventions to extracellular matrix maintenance and age-associated pathologies [75]. Such treatments may be based on the manipulation of ER signalling, as have previously been achieved of ERβ selective agonist effects on wound repair [52, 58, 76]. The therapeutic use of estrogen has become controversial in the post-Million Women and Women’s Health Initiative era [77, 78]. However, the skin offers a unique opportunity for topical estrogen treatment, avoiding many potential systemic side effects. Studies are now needed to determine the relative role of the estrogen receptors, ERα and ERβ in elastic fibre maintenance.

In conclusion we have shown that acute estrogen deprivation significantly alters the mechanical properties of the skin, leaving it both weaker and more lax. These structural changes are accompanied by a dramatic reduction in the volume of elastic fibres, correlated with significant subcutaneous adipose hypertrophy and increased gelatinase activity. Additionally we have demonstrated HDFs to be highly responsive to estrogen treatment up-regulating a number of ECM proteins especially those related to elastic fibre formation. Taken together these findings suggest a previously unappreciated role for dynamic elastic fibre changes in hormone-associated skin ageing. Future studies are now essential to elucidate the functional consequences for skin homeostasis and susceptibility to injury.

## Supporting information

Mass spectrometry data

## SUPPLEMENTARY DATA

Mass spectrometry data is available online.

## ACKNOWLEDGEMENTS

This research was supported by: an AgeUK Senior Fellowship (MJH), a Biotechnology and Biological Sciences Research Council (BBSRC) doctoral training programme (DTP) studentship (CRS); a BBSRC David Phillips Fellowship (BB/L024551/1) (JS). Mass spectrometry was carried out at the Wellcome Centre for Cell-Matrix Research (WCCMR; 203128/Z/16/Z) Proteomics Core Facility. We thank Drs. Ronan O-Cualain, Stacey Warwood, Julian Selley and David Knight for Core Facility support.

## References

[1] V.M. Tutino, M. Mandelbaum, A. Takahashi, L.C. Pope, A. Siddiqui, J. Kolega, H. Meng, Hypertension and estrogen deficiency augment aneurysmal remodeling in the rabbit circle of Willis in response to carotid ligation, Anat. Rec. 298 (2015) 1903–1910.

[2] M.K. Glassberg, R. Choi, V. Manzoli, S. Shahzeidi, P. Rauschkolb, R. Voswinckel, M. Aliniazee, X.M. Xia, S.J. Elliot, 17 beta-estradiol replacement reverses age-related lung disease in estrogen-deficient C57BL/6J mice, Endocrinology 155 (2014) 441–448.

[3] W. Most, L. Schot, A. Ederveen, L. Vanderweepals, S. Papapoulos, C. Lowik, In-vitro and ex-vivo evidence that estrogens suppress increased bone-resorption induced by ovariectomy or PTH stimulation through an effect on osteoclastogenesis J. Bone Miner. Res. 10 (1995) 1523–1530.

[4] K. Tsukahara, H. Nakagawa, S. Moriwaki, S. Kakuo, A. Ohuchi, Y. Takema, G. Imokawa, Ovariectomy is sufficient to accelerate spontaneous skin ageing and to stimulate ultraviolet irradiation-induced photoageing of murine skin, Br. J. Dermatol. 151 (2004) 984–994.

[5] M.J. Hardman, G.S. Ashcroft, Estrogen, not intrinsic aging, is the major regulator of delayed human wound healing in the elderly, Genome Biol. 9 (2008) 17.

[6] C. Escoffier, J. Derigal, A. Rochefort, R. Vasselet, J.L. Leveque, P.G. Agache, Age-related mechanical properties of human skin - an in vivo study, J. Invest. Dermatol. 93 (1989) 353–357.

[7] S.A. Thurstan, N.K. Gibbs, A.K. Langton, C.E.M. Griffiths, R.E.B. Watson, M.J. Sherratt, Chemical consequences of cutaneous photoageing, Chem. Cent. J. 6 (2012) 7.

[8] S. Verdier-Sevrain, F. Bonte, B. Gilchrest, Biology of estrogens in skin: implications for skin aging, Exp. Dermatol. 15 (2006) 83–94.

[9] M.A. Farage, K.W. Miller, E. Berardesca, H.I. Maibach, Clinical implications of aging skin cutaneous disorders in the elderly, Am. J. Clin. Dermatol. 10 (2009) 73–86.

[10] F.M. Hendriks, D. Brokken, C.W.J. Oomens, D.L. Bader, F.P.T. Baaijens, The relative contributions of different skin layers to the mechanical behavior of human skin in vivo using suction experiments, Med. Eng. Phys. 28 (2006) 259–266.

[11] O.S. Osman, J.L. Selway, P.E. Harikumar, C.J. Stocker, E.T. Wargent, M.A. Cawthorne, S. Jassim, K. Langlands, A novel method to assess collagen architecture in skin, BMC Bioinformatics 14 (2013) 10.

[12] H.K. Graham, N.W. Hodson, J.A. Hoyland, S.J. Millward-Sadler, D. Garrod, A. Scothern, C.E.M. Griffiths, R.E.B. Watson, T.R. Cox, J.T. Erler, A.W. Trafford, M.J. Sherratt, Tissue section AFM: In situ ultrastructural imaging of native biomolecules, Matrix Biol. 29 (2010) 254–260.

[13] M. Brincat, C.F. Moniz, S. Kabalan, E. Versi, T. Odowd, A.L. Magos, J. Montgomery, J.W.W. Studd, Decline in skin collagen content and metacarpal index after the menopause and its prevention with sex-hormone replacement, Br. J. Obstet. Gynaecol. 94 (1987) 126–129.

[14] A.V.D. Sauerbronn, A.M. Fonseca, V.R. Bagnoli, P.H. Saldiva, J.A. Pinotti, The effects of systemic hormonal replacement therapy on the skin of postmenopausal women, Int. J. Gynecol. Obstet. 68 (2000) 35–41.

[15] M. Brincat, E. Versi, C.F. Moniz, A. Magos, J. Detrafford, J.W.W. Studd, Skin collagen changes in postmenopausal women receiving different regimens of estrogen therapy, Obstet. Gynecol. 70 (1987) 123–127.

[16] C.R. Saville, V. Mallikarjun, D.F. Holmes, E. Emmerson, B. Derby, J. Swift, M.J. Sherratt, M.J. Hardman, Matrix proteomics and mechanics compared in ageing versus oestrogen-deficient skin, BioRxiv doi.org/10.1101/570481 (2019).

[17] H. Kafantari, E. Kounadi, M. Fatourous, M. Milonakis, M. Tzaphlidou, Structural alterations in rat skin and bone collagen fibrils induced by ovariectomy, Bone 26 (2000) 349–353.

[18] M. Fang, E.L. Goldstein, A.S. Turner, C.M. Les, B.G. Orr, G.J. Fisher, K.B. Welch, E.D. Rothman, M.M.B. Holl, Type I collagen D-spacing in fibril bundles of dermis, tendon, and bone: bridging between nano- and micro-level tissue hierarchy, ACS Nano 6 (2012) 9503–9514.

[19] M. Markiewicz, Y. Asano, S. Znoyko, Y. Gong, D.K. Watson, M. Trojanowska, Distinct effects of gonadectomy in mate and female mice on collagen fibrillogenesis in the skin, J. Dermatol. Sci. 47 (2007) 217–226.

[20] M. Brincat, C.J. Moniz, J.W.W. Studd, A. Darby, A. Magos, G. Emburey, E. Versi, Long-term effects of the menopause and sex-hormones on skin thickness, Br. J. Obstet. Gynaecol. 92 (1985) 256–259.

[21] Y.X. He, G. Zhang, X.H. Pan, Z. Liu, L.Z. Zheng, C.W. Chan, K.M. Lee, Y.P. Cao, G. Li, L. Wei, L.K. Hung, K.S. Leung, L. Qin, Impaired bone healing pattern in mice with ovariectomy-induced osteoporosis: A drill-hole defect model, Bone 48 (2011) 1388–1400.

[22] Y.H. Huang, Q.H. Zhang, Genistein reduced the neural apoptosis in the brain of ovariectomised rats by modulating mitochondrial oxidative stress, Br. J. Nutr. 104 (2010) 1297–1303.

[23] M.A. Cavasin, S.S. Sankey, A.L. Yu, S. Menon, X.P. Yang, Estrogen and testosterone have opposing effects on chronic cardiac remodeling and function in mice with myocardial infarction, Am. J. Physiol.-Heart Circul. Physiol. 284 (2003) H1560–H1569.

[24] M. Fang, K.G. Liroff, A.S. Turner, C.M. Les, B.G. Orr, M.M.B. Holl, Estrogen depletion results in nanoscale morphology changes in dermal collagen, J. Invest. Dermatol. 132 (2012) 1791–1797.

[25] K. Tsukahara, S. Moriwaki, A. Ohuchi, T. Fujimura, Y. Takema, Ovariectomy accelerates photoaging of rat skin, Photochem. Photobiol. 73 (2001) 525–531.

[26] J.M. Wallace, B. Erickson, C.M. Les, B.G. Orr, M.M.B. Holl, Distribution of type I collagen morphologies in bone: Relation to estrogen depletion, Bone 46 (2010) 1349–1354.

[27] F. Ramirez, D.B. Rifkin, Extracellular microfibrils: contextual platforms for TGF beta and BMP signaling, Curr. Opin. Cell Biol. 21 (2009) 616–622.

[28] G. Sengle, L.V. Sakai, The fibrillin microfibril scaffold: A niche for growth factors and mechanosensation?, Matrix Biol. 47 (2015) 3–12.

[29] C.M. Kielty, M.J. Sherratt, C.A. Shuttleworth, Elastic fibres, J. Cell Sci. 115 (2002) 2817–2828.

[30] G.M. Bressan, D. Dagagordini, A. Colombatti, I. Castellani, V. Marigo, D. Volpin, EMILIN, a component of elastic fibers preferentially located at the elastin-microfibrils interface, J. Cell Biol. 121 (1993) 201–212.

[31] M.A. Gibson, J.S. Kumaratilake, E.G. Cleary, The protein-components of the 12-nanometer microfibrils of elastic and non-elastic tissues, J. Biol. Chem. 264 (1989) 4590–4598.

[32] M.J. Sherratt, Tissue elasticity and the ageing elastic fibre, Age 31 (2009) 305–325.

[33] S.A. Hibbert, R.E.B. Watson, N.K. Gibbs, P. Costello, C. Baldock, A.S. Weiss, C.E.M. Griffiths, M.J. Sherratt, A potential role for endogenous proteins as sacrificial sunscreens and antioxidants in human tissues, Redox Biol. 5 (2015) 101–113.

[34] T. Ezure, S. Amano, Increment of subcutaneous adipose tissue is associated with decrease of elastic fibres in the dermal layer, Exp. Dermatol. 24 (2015) 924–929.

[35] M. Just, M. Ribera, E. Monso, J.C. Lorenzo, C. Ferrandiz, Effect of smoking on skin elastic fibres: morphometric and immunohistochemical analysis, Br. J. Dermatol. 156 (2007) 85–91.

[36] P. Newsholme, V.F. Cruzat, K.N. Keane, R. Carlessi, P.I.H. de Bittencourt, Molecular mechanisms of ROS production and oxidative stress in diabetes, Biochem. J. 473 (2016) 4527–4550.

[37] R. Akhtar, J.K. Cruickshank, X. Zhao, L.A. Walton, N.J. Gardiner, S.D. Barrett, H.K. Graham, B. Derby, M.J. Sherratt, Localized micro- and nano-scale remodelling in the diabetic aorta, Acta Biomater. 10 (2014) 4843–4851.

[38] I. Baeza, J. Fdez-Tresguerres, C. Ariznavarreta, M. De la Fuente, Effects of growth hormone, melatonin, oestrogens and phytoestrogens on the oxidized glutathione (GSSG)/reduced glutathione (GSH) ratio and lipid peroxidation in aged ovariectomized rats, Biogerontology 11 (2010) 687–701.

[39] G. Bottai, R. Mancina, M. Muratori, P. Di Gennaro, T. Lotti, 17 beta-estradiol protects human skin fibroblasts and keratinocytes against oxidative damage, J. Eur. Acad. Dermatol. Venereol. 27 (2013) 1236–1243.

[40] F. Lizcano, G. Guzman, Estrogen deficiency and the origin of obesity during menopause, Biomed Res. Int. (2014) 11.

[41] E. Emmerson, L. Campbell, F.C.J. Davies, N.L. Ross, G.S. Ashcroft, A. Krust, P. Chambon, M.J. Hardman, Insulin-like growth factor-1 promotes wound healing in estrogen-deprived mice: New insights into cutaneous IGF-1R/ER alpha cross talk, J. Invest. Dermatol. 132 (2012) 2838–2848.

[42] N.H. Cooper, J.P. Balachandra, M.J. Hardman, Global gene expression analysis in PKC alpha(-/-) mouse skin reveals structural changes in the dermis and defective wound granulation tissue, J. Invest. Dermatol. 135 (2015) 3173–3182.

[43] G. Gomori, Aldehyde-fuchsin - a new stain for elastic tissue, Am. J. Clin. Pathol. 20 (1950) 665–666.

[44] Y.T. Kim, J.C. Park, S.H. Choi, K.S. Cho, G.I. Im, B.S. Kim, C.S. Kim, The dynamic healing profile of human periodontal ligament stem cells: histological and immunohistochemical analysis using an ectopic transplantation model, J. Periodont. Res. 47 (2012) 514–524.

[45] J.C. McConnell, O.V. O’Connell, K. Brennan, L. Weiping, M. Howe, L. Joseph, D. Knight, R. O’Cualain, Y. Lim, A. Leek, R. Waddington, J. Rogan, S.M. Astley, A. Gandhi, C.C. Kirwan, M.J. Sherratt, C.H. Streuli, Increased peri-ductal collagen micro-organization may contribute to raised mammographic density, Breast Cancer Res. 18 (2016) 17.

[46] H.K. Graham, R. Akhtar, C. Kridiotis, B. Derby, T. Kundu, A.W. Trafford, M.J. Sherratt, Localised micro-mechanical stiffening in the ageing aorta, Mech. Ageing. Dev. 132 (2011) 459–467.

[47] R. Rezakhaniha, A. Agianniotis, J.T.C. Schrauwen, A. Griffa, D. Sage, C.V.C. Bouten, F.N. van de Vosse, M. Unser, N. Stergiopulos, Experimental investigation of collagen waviness and orientation in the arterial adventitia using confocal laser scanning microscopy, Biomech. Model. Mechanobiol. 11 (2012) 461–473.

[48] C.M. Lapiere, B. Nusgens, G.E. Pierard, Interaction between collagen type-1 and type-3 in conditioning bundles organization, Connect. Tissue Res. 5 (1977) 21–29.

[49] P. Lu, K. Takai, V.M. Weaver, Z. Werb, Extracellular matrix degradation and remodeling in development and disease, Cold Spring Harb. Perspect. Biol. 3 (2011).

[50] O.R.F. Mook, C. Van Overbeek, E.G. Ackema, F. Van Maldegem, W.M. Frederiks, In situ localization of gelatinolytic activity in the extracellular matrix of metastases of colon cancer in rat liver using quenched fluorogenic DQ-gelatin, J. Histochem. Cytochem. 51 (2003) 821–829.

[51] N. Philips, J. Devaney, Beneficial regulation of type I collagen and matrixmetalloproteinase-1 expression by estrogen, progesterone, and its combination in skin fibroblasts, J. Am. Aging Assoc. 26 (2003) 59–62.

[52] L. Campbell, E. Emmerson, F. Davies, S.C. Gilliver, A. Krust, P. Chambon, G.S. Ashcroft, M.J. Hardman, Estrogen promotes cutaneous wound healing via estrogen receptor beta independent of its antiinflammatory activities, J. Exp. Med. 207 (2010) 1825–1833.

[53] F. Polito, H. Marini, A. Bitto, N. Irrera, M. Vaccaro, E.B. Adamo, A. Micali, F. Squadrito, L. Minutoli, D. Altavilla, Genistein aglycone, a soy-derived isoflavone, improves skin changes induced by ovariectomy in rats, Br. J. Pharmacol. 165 (2012) 994–1005.

[54] G.E. Pierard, C. Letawe, A. Dowlati, C. Pierardfranchimont, Effect of hormone replacement therapy for menopause on the mechanical-properties of skin, J. Am. Geriatr. Soc. 43 (1995) 662–665.

[55] P.G. Agache, C. Monneur, J.L. Leveque, J. Derigal, Mechanical properties and Young’s modulus of human skin in vivo, Arch. Dermatol. Res. 269 (1980) 221–232.

[56] G. Boyer, L. Laquieze, A. Le Bot, S. Laquieze, H. Zahouani, Dynamic indentation on human skin in vivo: ageing effects, Skin Res. Technol. 15 (2009) 55–67.

[57] I. Levental, P.C. Georges, P.A. Janmey, Soft biological materials and their impact on cell function, Soft Matter 3 (2007) 299–306.

[58] M.J. Hardman, E. Emmerson, L. Campbell, G.S. Ashcroft, Selective estrogen receptor modulators accelerate cutaneous wound healing in ovariectomized female mice, Endocrinology 149 (2008) 551–557.

[59] M. El-Domyati, S. Attia, F. Saleh, D. Brown, D.E. Birk, F. Gasparro, H. Ahmad, J. Uitto, Intrinsic aging vs. photoaging: a comparative histopathological, immunohistochemical, and ultrastructural study of skin, Exp. Dermatol. 11 (2002) 398–405.

[60] H. Oxlund, Relationships between the biomechanical properties, composition and molecular-structure of connective tissues, Connect. Tissue Res. 15 (1986) 65–72.

[61] G.S. Ashcroft, C.M. Kielty, M.A. Horan, M.W.J. Ferguson, Age-related changes in the temporal and spatial distributions of fibrillin and elastin mRNAs and proteins in acute cutaneous wounds of healthy humans, J. Pathol. 183 (1997) 80–89.

[62] J.M. Davidson, K.E. Hill, J.L. Alford, Developmental-changes in collagen and elastin biosynthesis in the porcine aorta, Dev. Biol. 118 (1986) 103–111.

[63] S. Chakraborti, M. Mandal, S. Das, A. Mandal, T. Chakraborti, Regulation of matrix metalloproteinases: An overview, Mol. Cell. Biochem. 253 (2003) 269–285.

[64] M. Matsumoto, A. Ibuki, T. Minematsu, J. Sugama, M. Horii, K. Ogai, T. Nishizawa, M. Dai, A. Sato, Y. Fujimoto, M. Okuwa, G. Nakagami, T. Nakatani, H. Sanada, Structural changes in dermal collagen and oxidative stress levels in the skin of Japanese overweight males, Int. J. Cosmetic Sci. 36 (2014) 477–484.

[65] R.E.B. Watson, N.K. Gibbs, C.E.M. Griffiths, M.J. Sherratt, Damage to skin extracellular matrix induced by UV exposure, Antioxid. Redox. Sign. 21 (2014) 1063–+.

[66] E. Emmerson, M.J. Hardman, The role of estrogen deficiency in skin ageing and wound healing, Biogerontology 13 (2012) 3–20.

[67] R. Doliana, S. Bot, G. Mungiguerra, A. Canton, S.P. Cilli, A. Colombatti, Isolation and characterization of EMILIN-2, a new component of the growing EMILINs family and a member of the EMI domain-containing superfamily, J. Biol. Chem. 276 (2001) 12003–12011.

[68] S. Bot, E. Andreuzzi, A. Capuano, A. Schiavinato, A. Colombatti, R. Doliana, Multiple-interactions among EMILIN1 and EMILIN2 N- and C-terminal domains, Matrix Biol. 41 (2015) 44–55.

[69] D.P. Reinhardt, T. Sasaki, B.J. Dzamba, D.R. Keene, M.L. Chu, W. Gohring, R. Timpl, L.Y. Sakai, Fibrillin-1 and fibulin-2 interact and are colocalized in some tissues, J. Biol. Chem. 271 (1996) 19489–19496.

[70] E.D. Son, J.Y. Lee, S. Lee, M.S. Kim, B.G. Lee, I.S. Chang, J.H. Chung, Topical application of 17 beta-estradiol increases extracellular matrix protein synthesis by stimulating TGF-beta signaling in aged human skin in vivo, J. Invest. Dermatol. 124 (2005) 1149–1161.

[71] R. Punnonen, P. Vaajalahti, K. Teisala, Local estriol treatment improves the structure of elastic fibers in the skin of postmenopausal women, Ann. Chir. Gynaecol. 76 (1987) 39–41.

[72] N. Heldring, A. Pike, S. Andersson, J. Matthews, G. Cheng, J. Hartman, M. Tujague, A. Strom, E. Treuter, M. Warner, J.A. Gustafsson, Estrogen receptors: How do they signal and what are their targets, Physiol. Rev. 87 (2007) 905–931.

[73] S.M. Aronica, B.S. Katzenellenbogen, Stimulation of estrogen receptor-mediated transcription and alteration in the phosphorylation state of the rat uterine estrogen-receptor by estrogen, cyclic adenosine-monophosphate, and insulin-like growth factor-I, Mol. Endocrinol. 7 (1993) 743–752.

[74] G. Hall, T.J. Phillips, Estrogen and skin: The effects of estrogen, menopause, and hormone replacement therapy on the skin, J. Am. Acad. Dermatol. 53 (2005) 555–568.

[75] V. Mallikarjun, J. Swift, Therapeutic manipulation of ageing: repurposing old dogs and discovering new tricks, EBioMedicine 14 (2016) 24–31.

[76] E. Emmerson, L. Campbell, G.S. Ashcroft, M.J. Hardman, Unique and synergistic roles for 17 beta-estradiol and macrophage migration inhibitory factor during cutaneous wound closure are cell type specific, Endocrinology 150 (2009) 2749–2757.

[77] E. Banks, V. Beral, D. Bull, G. Reeves, J. Austoker, R. English, J. Patnick, R. Peto, M. Vessey, M. Wallis, S. Abbott, E. Bailey, K. Baker, A. Balkwill, I. Barnes, J. Black, A. Brown, B. Cameron, K. Canfell, A. Cliff, B. Crossley, E. Couto, S. Davies, D. Ewart, S. Ewart, D. Ford, L. Gerrard, A. Goodill, J. Green, W. Gray, E. Hilton, A. Hogg, J. Hooley, A. Hurst, S.W. Kan, C. Keene, N. Langston, A. Roddam, P. Saunders, E. Sherman, M. Simmonds, E. Spencer, H. Strange, A. Timadjer, C. Million Women Study, Breast cancer and hormone-replacement therapy in the Million Women Study, Lancet 362 (2003) 419–427.

[78] V. Beral, D. Bull, J. Green, G. Reeves, C. Million Women Study, Ovarian cancer and hormone replacement therapy in the Million Women Study, Lancet 369 (2007) 1703–1710.

